# Solubility-Weighted Index: fast and accurate prediction of protein solubility

**DOI:** 10.1101/2020.02.15.951012

**Authors:** Bikash K. Bhandari, Paul P. Gardner, Chun Shen Lim

## Abstract

**Motivation:** Recombinant protein production is a widely used technique in the biotechnology and biomedical industries, yet only a quarter of target proteins are soluble and can therefore be purified.

**Results:** We have discovered that global structural flexibility, which can be modeled by normalised B-factors, accurately predicts the solubility of 12,216 recombinant proteins expressed in *Escherichia coli*. We have optimised B-factors, and derived a new set of values for solubility scoring that further improves prediction accuracy. We call this new predictor the ‘Solubility-Weighted Index’ (SWI). Importantly, SWI outperforms many existing protein solubility prediction tools. Furthermore, we have developed ‘SoDoPE’ (Soluble Domain for Protein Expression), a web interface that allows users to choose a protein region of interest for predicting and maximising both protein expression and solubility.

**Availability:** The SoDoPE web server and source code are freely available at https://tisigner.com/sodope and https://github.com/Gardner-BinfLab/TISIGNER-ReactJS, respectively. The code and data for reproducing our analysis can be found at https://github.com/Gardner-BinfLab/SoDoPE_paper2020.

## INTRODUCTION

High levels of protein expression and solubility are two major requirements of successful recombinant protein production (Esposito and Chatterjee 2006). However, recombinant protein production is a challenging process. Almost half of recombinant proteins fail to be expressed and half of the successfully expressed proteins are insoluble (http://targetdb.rcsb.org/metrics/). These failures hamper protein research, with particular implications for structural, functional and pharmaceutical studies that require soluble and concentrated protein solutions (Kramer et al. 2012; Hou et al. 2018). Therefore, solubility prediction and protein engineering for enhanced solubility is an active area of research. Notable protein engineering approaches include mutagenesis, truncation (i.e., expression of partial protein sequences), or fusion with a solubility-enhancing tag (Waldo 2003; Esposito and Chatterjee 2006; Trevino, Martin Scholtz, and Nick Pace 2007; Chan et al. 2010; Kramer et al. 2012; Costa et al. 2014).

Protein solubility, at least in part, depends upon extrinsic factors such as ionic strength, temperature and pH, as well as intrinsic factors—the physicochemical properties of the protein sequence and structure, including molecular weight, amino acid composition, hydrophobicity, aromaticity, isoelectric point, structural propensities and the polarity of surface residues (Wilkinson and Harrison 1991; Chiti et al. 2003; Tartaglia et al. 2004; Diaz et al. 2010). Many solubility prediction tools have been developed around these features using statistical models (e.g., linear and logistic regression) or other machine learning models (e.g., support vector machines and neural networks) (Hirose and Noguchi 2013; Habibi et al. 2014; Hebditch et al. 2017; Sormanni et al. 2017; Heckmann et al. 2018; Z. Wu et al. 2019; Yang, Wu, and Arnold 2019).

In this study, we investigated the experimental outcomes of 12,216 recombinant proteins expressed in *Escherichia coli* from the ‘Protein Structure Initiative:Biology’ (PSI:Biology) (Chen et al. 2004; Acton et al. 2005). We showed that protein structural flexibility is more accurate than other protein sequence properties in predicting solubility (Craveur et al. 2015; M. Vihinen, Torkkila, and Riikonen 1994). Flexibility is a standard feature that appears to have been overlooked in previous solubility prediction attempts. On this basis, we derived a set of 20 values for the standard amino acid residues and used them to predict solubility. We call this new predictor the ‘Solubility-Weighted Index’ (SWI). SWI is a powerful predictor of solubility, and a good proxy for global structural flexibility. In addition, SWI outperforms many existing *de novo* protein solubility prediction tools.

## RESULTS

### Global structural flexibility performs well at predicting protein solubility

We sought to understand what makes a protein soluble, and develop a fast and accurate approach for solubility prediction. To determine which protein sequence properties accurately predict protein solubility, we analysed 12,216 target proteins from over 196 species that were expressed in *E. coli* (the PSI:Biology dataset; see Supplementary Fig S1 and Table S1A) (Chen et al. 2004; Acton et al. 2005). These proteins were expressed either with a C-terminal or N-terminal polyhistidine fusion tag (pET21_NESG and pET15_NESG expression vectors, N=8,780 and 3,436, respectively). They were previously curated and labeled as ‘Protein_Soluble’ or ‘Tested_Not_Soluble’ (Seiler et al. 2014), based on the soluble analysis of cell lysate using SDS-PAGE (R. Xiao et al. 2010). A total of 8,238 recombinant proteins were found to be soluble, in which 6,432 of them belong to the pET21_NESG dataset. Both the expression system and solubility analysis method are commonly used (Costa et al. 2014). Therefore, this collection of data captures a broad range of protein solubility issues.

We evaluated nine standard and 9,920 miscellaneous protein sequence properties using the Biopython’s ProtParam module and ‘protr’ R package, respectively (Cock et al. 2009; N. Xiao et al. 2015). For example, the standard properties include the Grand Average of Hydropathy (GRAVY), secondary structure propensities, protein structural flexibility etc., whereas miscellaneous properties include amino acid composition, autocorrelation, etc. Strikingly, protein structural flexibility outperformed other features in solubility prediction [Area Under the ROC Curve (AUC) = 0.67; Fig 1, Supplementary Fig S2 and Table S2].

**Fig 1.**
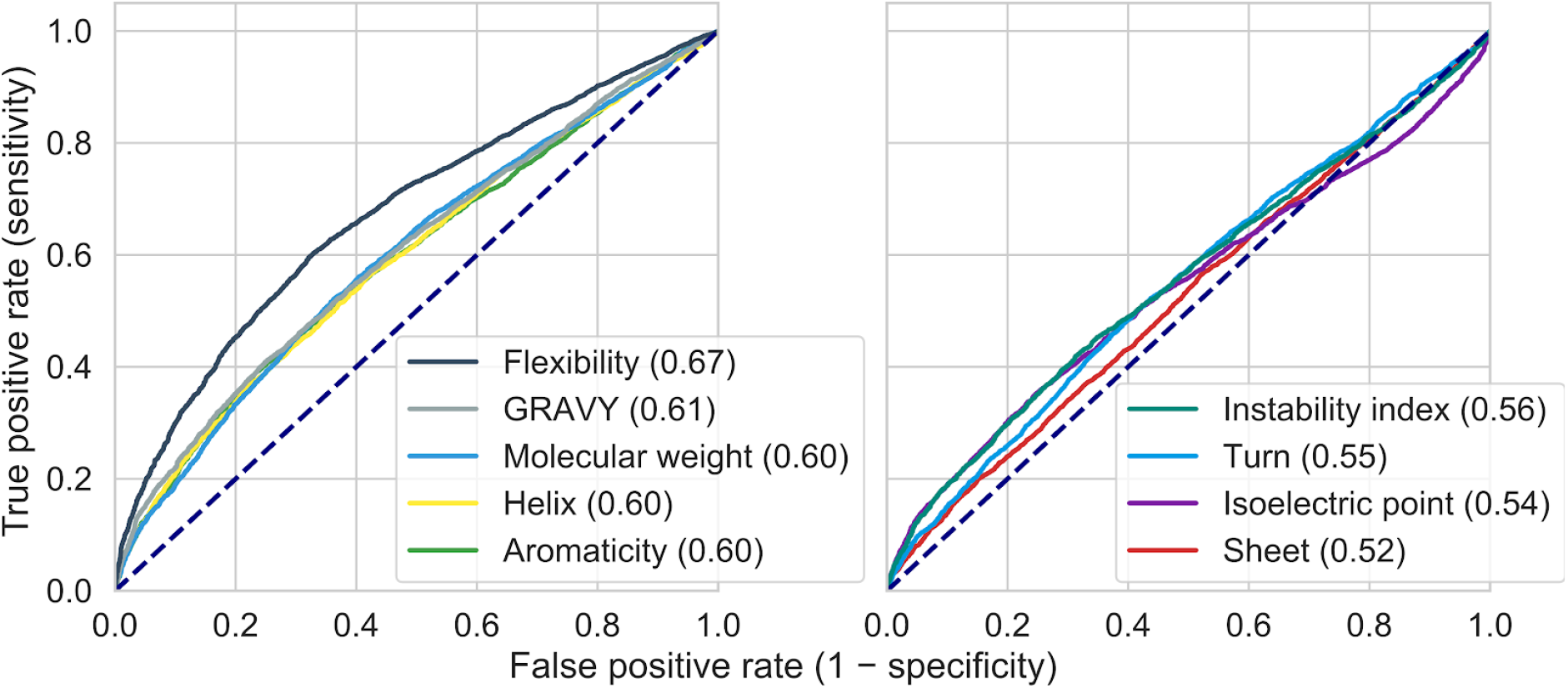
Global structural flexibility outperforms the other standard protein sequence properties in protein solubility prediction. ROC analysis of the standard protein sequence features for predicting the solubility of 12,216 recombinant proteins expressed in *E. coli* (the PSI:Biology dataset). AUC scores (perfect = 1.00, random = 0.50) are shown in parentheses. The ROC curves are shown in two separate panels for clarity. Dashed lines denote the performance of random classifiers. See also Supplementary Fig S2 and Table S2. AUC, Area Under the ROC Curve; GRAVY, Grand Average of Hydropathy; PSI:Biology, Protein Structure Initiative:Biology; ROC, Receiver Operating Characteristic.

### The Solubility-Weighted Index (SWI) is an improved predictor of solubility

Protein structural flexibility, in particular, the flexibility of local regions, is often associated with function (Craveur et al. 2015). The calculation of flexibility is usually performed by assigning a set of 20 normalised B-factors—a measure of vibration of C-alpha atoms (see Supplementary Notes)—to a protein sequence and averaging the values by a sliding window approach (Ragone et al. 1989; Karplus and Schulz 1985; M. Vihinen, Torkkila, and Riikonen 1994; Smith et al. 2003). We reasoned that such sliding window approach can be approximated by a more straightforward arithmetic mean for calculating global structural flexibility (see Supplementary Notes). We determined the correlation between flexibility (Vihinen *et al.’* s sliding window approach as implemented in Biopython) and solubility scores calculated as follows:

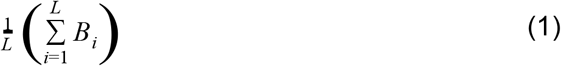

where *B_i_* is the normalised B-factor of the amino acid residue at the position *i*, and *L* is the sequence length. We obtained a strong correlation for the PSI:Biology dataset (Spearman’s rho = 0.98, P-value below machine’s underflow level). Therefore, we reasoned that the sliding window approach is not necessary for our purpose.

We applied this arithmetic mean approach (i.e., sequence composition scoring) to the PSI:Biology dataset and compared four sets of previously published, normalised B-factors (Bhaskaran and Ponnuswamy 1988; Ragone et al. 1989; M. Vihinen, Torkkila, and Riikonen 1994; Smith et al. 2003) Among these sets of B-factors, sequence composition scoring using the most recently published set of normalised B-factors produced the highest AUC score (Supplementary Fig S3, AUC = 0.66).

To improve the prediction accuracy of solubility, we iteratively refined the weights of amino acid residues using the Nelder-Mead optimisation algorithm (Nelder and Mead 1965). To avoid testing and training on similar sequences, we generated 10 cross-validation sets with a maximised heterogeneity between these subsets (i.e. no similar sequences between subsets). We first clustered all 12,216 PSI:Biology protein sequences using a 40% similarity threshold using USEARCH to produce 5,050 clusters with remote similarity (see Methods and Supplementary Fig S4). The clusters were grouped into 10 cross-validation sets of approximately 1,200 sequences each manually. We did not select a representative sequence for each cluster as about 12% of clusters contain a mix of soluble and insoluble proteins (Supplementary Fig S4C). More importantly, to address the issues of sequence similarity and imbalanced classes, we performed 1,000 bootstrap resamplings for each cross-validation step (Fig 2A and Supplementary Fig S5). We calculated the solubility scores using the optimised weights as Equation 1 and the AUC scores for each cross-validation step. Our training and test AUC scores were 0.72 ± 0.00 and 0.71 ± 0.01, respectively, showing an improvement over flexibility in solubility prediction (mean ± standard deviation; Fig 2B and Supplementary Table S3).

**Fig 2.**
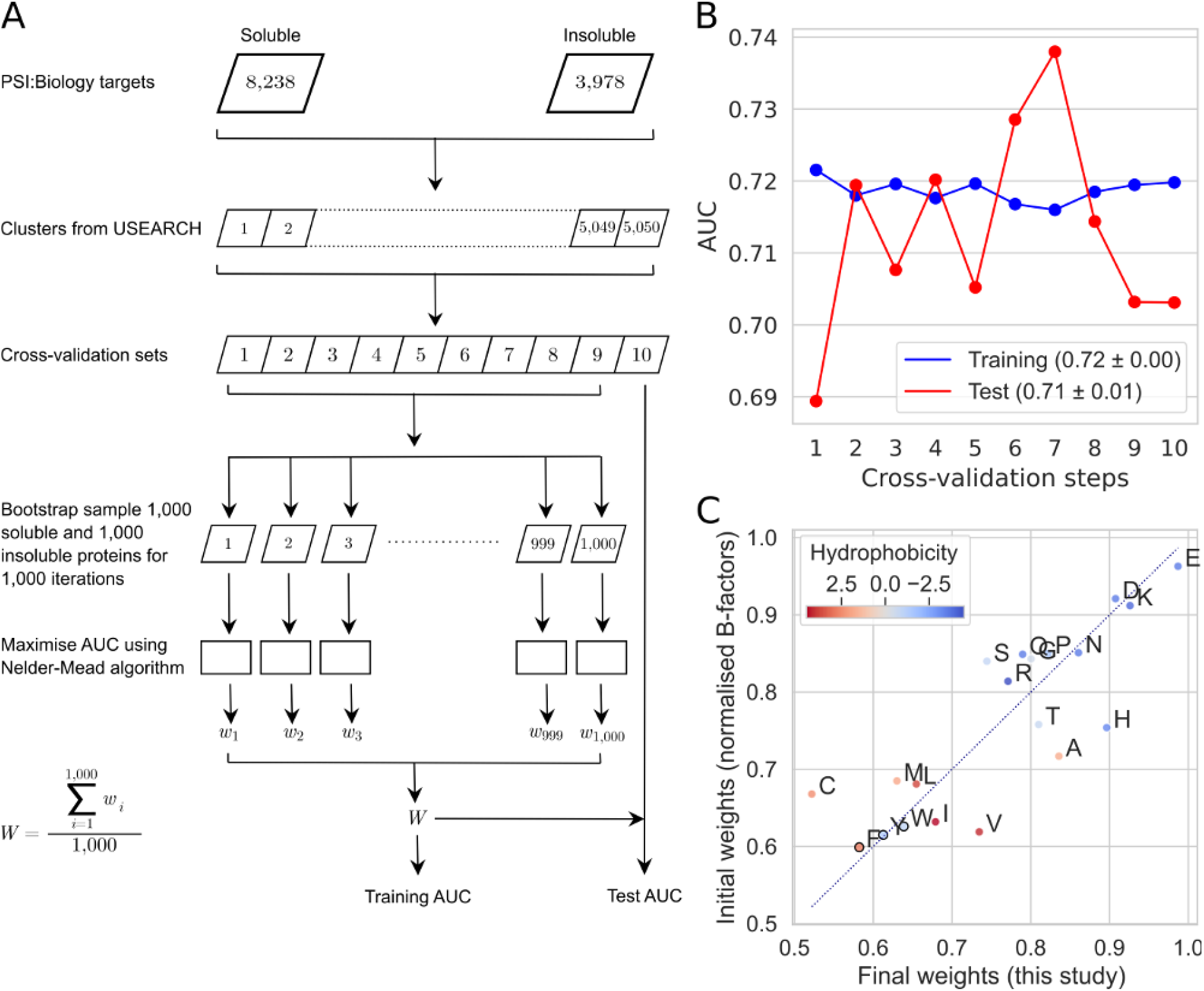
Derivation of the Solubility-Weighted Index (SWI). **(A)** Flow chart shows an iterative refinement of the most recently published set of normalised B-factors for solubility prediction (Smith et al. 2003). The solubility score of a protein sequence was calculated using a sequence composition scoring approach (Equation 1, using optimised weights *W*, instead of normalised B-factors *B*). These scores were used to compute the AUC scores for training and test datasets. **(B)** Training and test performance of solubility prediction using optimised weights for 20 amino acid residues in a 10-fold cross-validation (mean AUC ± standard deviation). Related data and figures are available as Supplementary Table S3 and Supplementary Fig S4 and S5. **(C)** Comparison between the 20 initial and final weights for amino acid residues. The final weights are derived from the arithmetic mean of the optimised weights from cross-validation. These weights are used to calculate SWI, the solubility score of a protein sequence, in the subsequent analyses. Filled circles, which represent amino acid residues, are colored by hydrophobicity (Kyte and Doolittle 1982). Solid black circles denote aromatic amino acid residues phenylalanine (F), tyrosine (Y), tryptophan (W). Dotted diagonal line represents no change in weight. See also Supplementary Table S4 and Fig S4. AUC, Area Under the ROC Curve; ROC, Receiver Operating Characteristic; *W*, arithmetic mean of the weights of an amino acid residue optimised from 1,000 bootstrap samples in a cross-validation step.

The final weights were derived from the arithmetic means of the weights for individual amino acid residues obtained cross-validation (Supplementary Table S4). We observed over a 20% change on the weights for cysteine (C) and histidine (H) residues (Fig 2C and Supplementary Table S4). These results are in agreement with the contributions of cysteine and histidine residues as shown in Supplementary Fig S2B. We call the solubility score of a protein sequence calculated using the final weights the Solubility-Weighted Index (SWI).

To validate the cross-validation results, we used a dataset independent of the PSI:Biology data known as eSOL (Niwa et al. 2009). This dataset consists of the solubility percentages of *E. coli* proteins determined using an *E. coli* cell-free system (N = 3,198). Our solubility scoring using the final weights showed a significant improved correlation with *E. coli* protein solubility over the initial weights (Smith *et al.’* s normalised B-factors) [Spearman’s rho of 0.50 (P = 9.46 × 10^-206^) versus 0.40 (P = 4.57 × 10^-120^)]. We repeated the correlation analysis by removing extra amino acid residues including His-tags from the eSOL sequences (MRGSHHHHHHTDPALRA and GLCGR at the N- and C-termini, respectively). This artificial dataset was created based on the assumption that His-tags have little effect on solubility. We observed a slight decrease in correlation for this artificial dataset (Spearman’s rho = 0.47, P= 3.67 × 10-176), which may be due to the effects of His-tag in solubility and/or the limitation(s) of our approach that may overfit to His-tag fusion proteins.

We performed Spearman’s correlation analysis for both the PSI:Biology and eSOL datasets. SWI shows the strongest correlation with solubility compared to the standard and 9,920 protein sequence properties (Fig 3 and Supplementary Fig S2, respectively). SWI also strongly correlates with flexibility, suggesting that SWI is also a good proxy for global structural flexibility.

**Fig 3.**
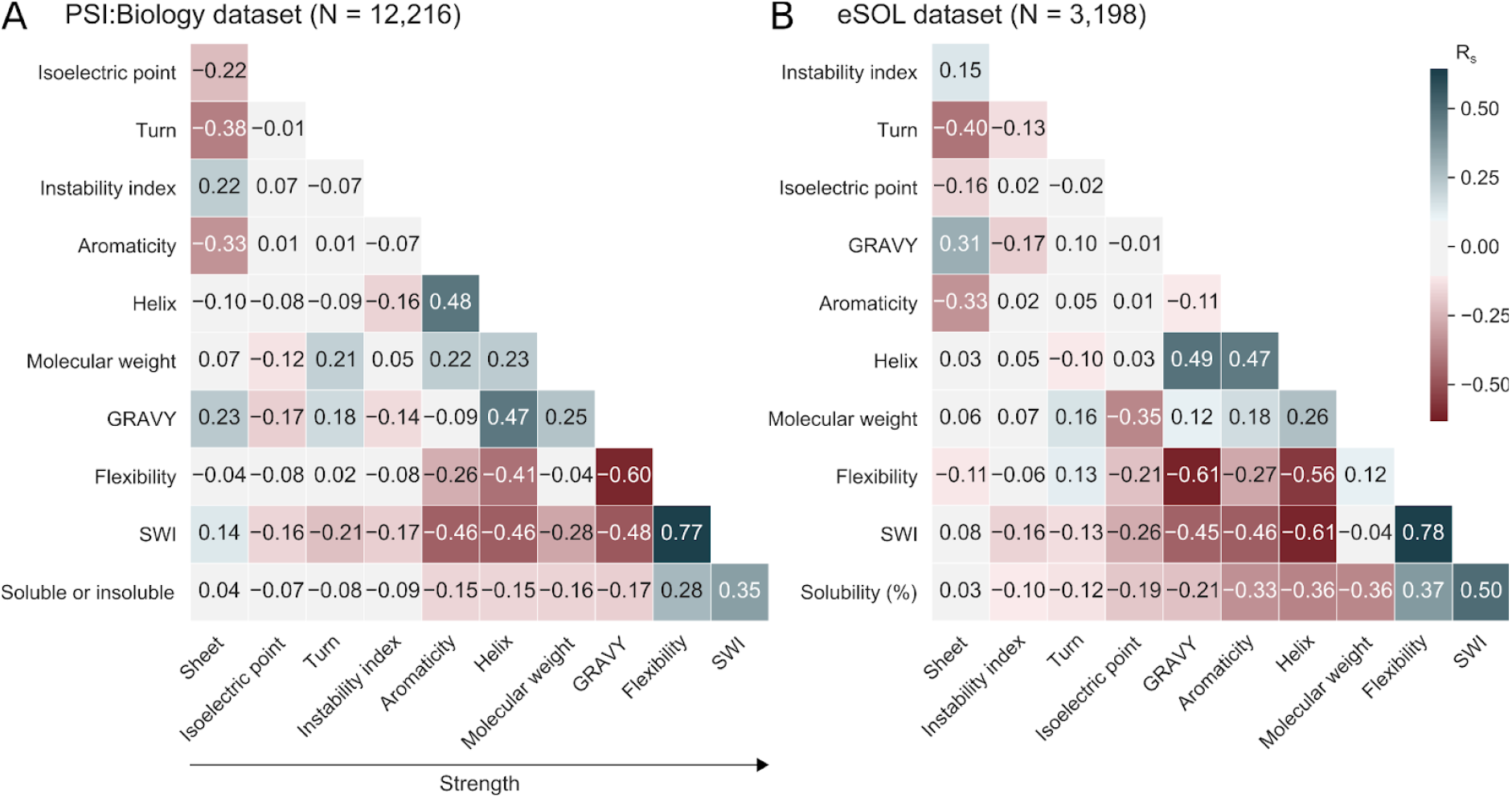
SWI strongly correlates with solubility. **(A)** Correlation matrix plot of the solubility of recombinant proteins expressed in *E. coli* and their standard protein sequence properties and SWI. These recombinant proteins are the PSI:Biology targets (N = 12,216) with a binary solubility status of ‘Protein_Soluble’ or ‘Tested_Not_Soluble’. Related data is available as Supplementary Table S5. **(B)** Correlation matrix plot of the solubility percentages of *E. coli* proteins and their standard protein sequence properties and SWI. The solubility percentages were previously determined using an *E. coli* cell-free system (eSOL, N = 3,198). Related data is available as Supplementary Table S6. GRAVY, Grand Average of Hydropathy;

PSI:Biology, Protein Structure Initiative:Biology; R_s_, Spearman’s rho; SWI, Solubility-Weighted Index.

We asked whether protein solubility can be predicted by surface amino acid residues. To address this question, we examined a previously published dataset for the protein surface ‘stickiness’ of 397 *E. coli* proteins (Levy, De, and Teichmann 2012). This dataset has the annotation for surface residues based on previously solved protein crystal structures. We observed little correlation between the protein surface ‘stickiness’ and the solubility data from eSOL (Spearman’s rho = 0.05, P = 0.34, N = 348; Supplementary Fig S6A). Next, we evaluated if amino acid composition scoring using surface residues is sufficient, optimising only the weights of surface residues should achieve similar or better results than SWI. As above, we iteratively refined the weights of surface residues using the Nelder-Mead optimisation algorithm. The method was initialised with Smith *et al.’* s normalised B-factors and a maximised correlation coefficient was the target. However, a low correlation was obtained upon convergence (Spearman’s rho = 0.18, P = 7.20 × 10^-4^; Supplementary Fig S6B). In contrast, the SWI of the full-length sequences has a much stronger correlation with solubility (Spearman’s rho = 0.46, P = 2.97 × 10^-19^; Supplementary Fig S6C). These results suggest that the full-length of sequences contributes to protein solubility, not just surface residues, in which solubility is modulated by cotranslational folding (Natan et al. 2018).

To understand the properties of soluble and insoluble proteins, we determined the enrichment of amino acid residues in the PSI:Biology targets relative to the eSOL sequences (see Methods). We observed that the PSI:Biology targets are enriched in charged residues lysine (K), glutamate (E) and aspartate (D), and depleted in aromatic residues tryptophan (W), albeit to a lesser extend for insoluble proteins (Supplementary Fig S7A). As expected, cysteine residues (C) are enriched in the PSI:Biology insoluble proteins, supporting previous findings that cysteine residues contribute to poor solubility in the *E. coli* expression system (Diaz et al. 2010; Wilkinson and Harrison 1991).

In addition, we compared the SWI of random sequences with the PSI:Biology and eSOL sequences. We included an analysis of random sequences to confirm whether SWI can distinguish between biological and random sequences. We found that the SWI scores of soluble proteins are higher than those of insoluble proteins (Supplementary Fig S7B), and that true biological sequences also tend to have higher SWI scores than random sequences, highlighting a potential evolutionary selection for solubility.

### SWI outperforms many protein solubility prediction tools

To confirm the usefulness of SWI in solubility prediction, we compared it with the existing tools Protein-Sol (Hebditch et al. 2017), CamSol v2.1 (Sormanni, Aprile, and Vendruscolo 2015; Sormanni et al. 2017), PaRSnIP (Rawi et al. 2018), DeepSol v0.3 (Khurana et al. 2018), the Wilkinson-Harrison model (Davis et al. 1999; Harrison 2000; Wilkinson and Harrison 1991), and ccSOL omics (Agostini et al. 2014). We did not include the specialised tools that model protein structural information such as surface geometry, surface charges and solvent accessibility because these tools require prior knowledge of protein tertiary structure. For example, Aggrescan3D and SOLart accept only PDB files that can be downloaded from the Protein Data Bank or produced using a homology modeling program (Kuriata et al. 2019; Hou et al. 2019). SWI outperforms other tools except for Protein-Sol in predicting *E. coli* protein solubility (Table 1, Fig 4A). Our SWI C program is also the fastest solubility prediction algorithm (Table 1, Fig 4B and Supplementary Table S7).

**Table 1.**
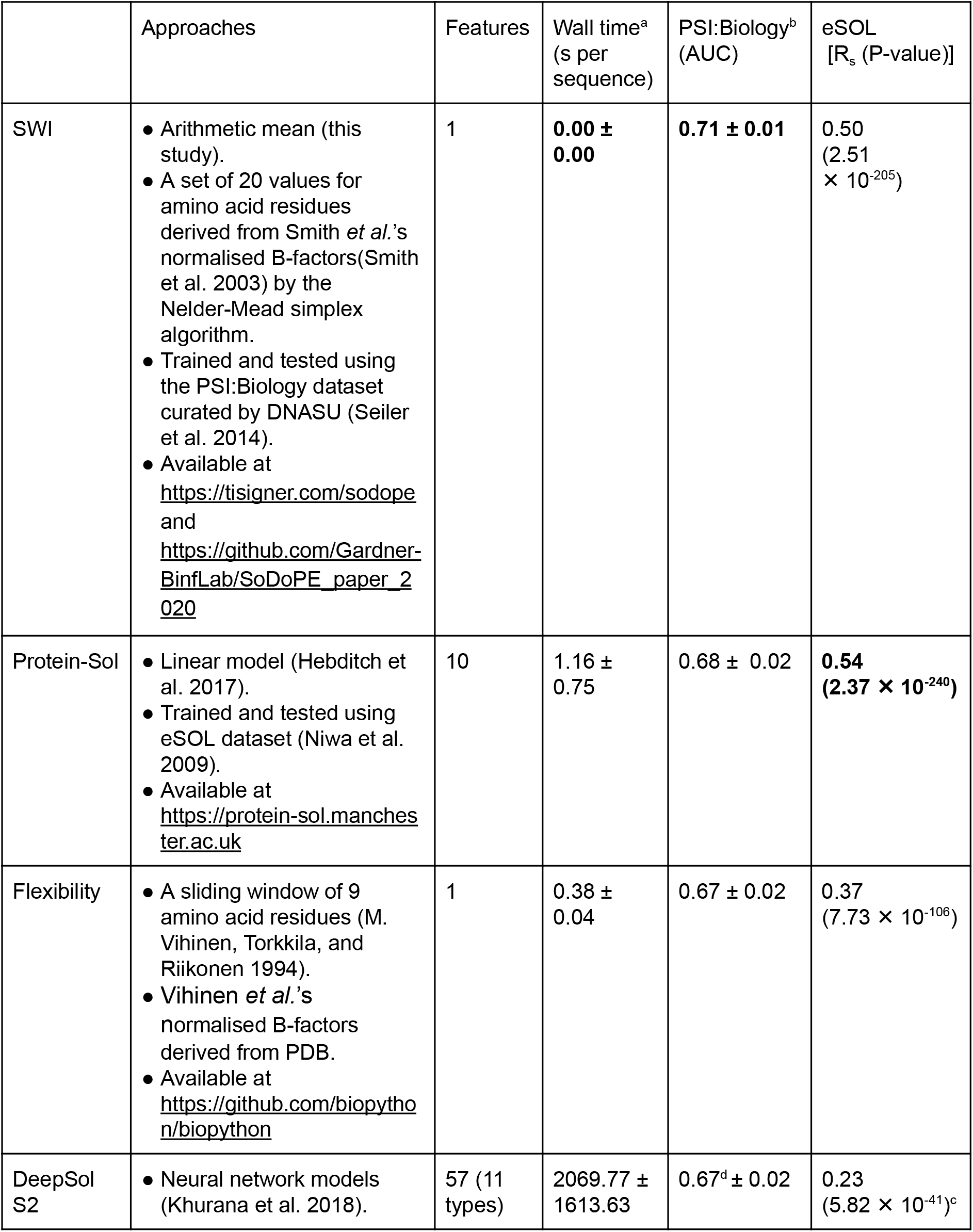
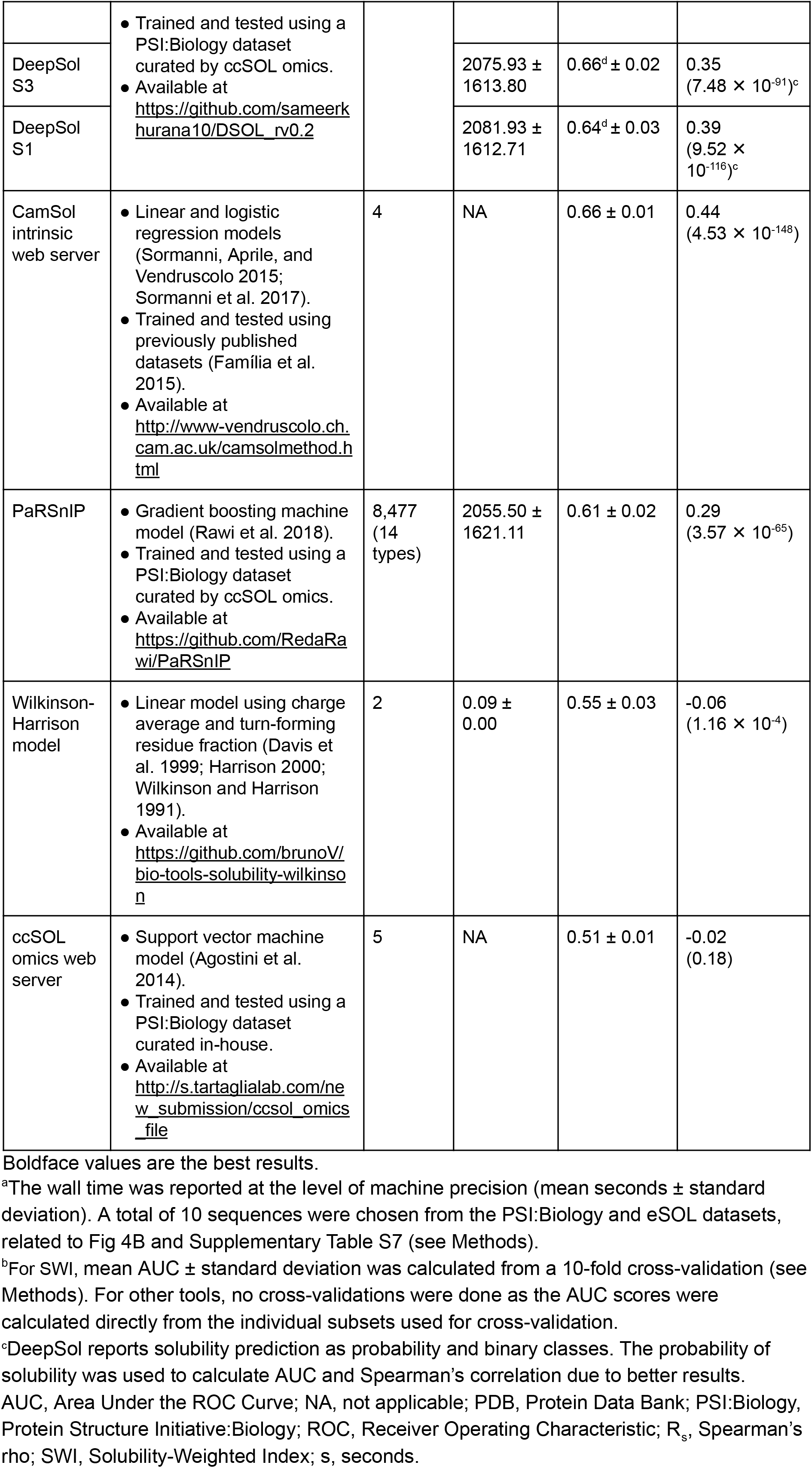
Comparison of protein solubility prediction methods and software.

**Fig 4.**
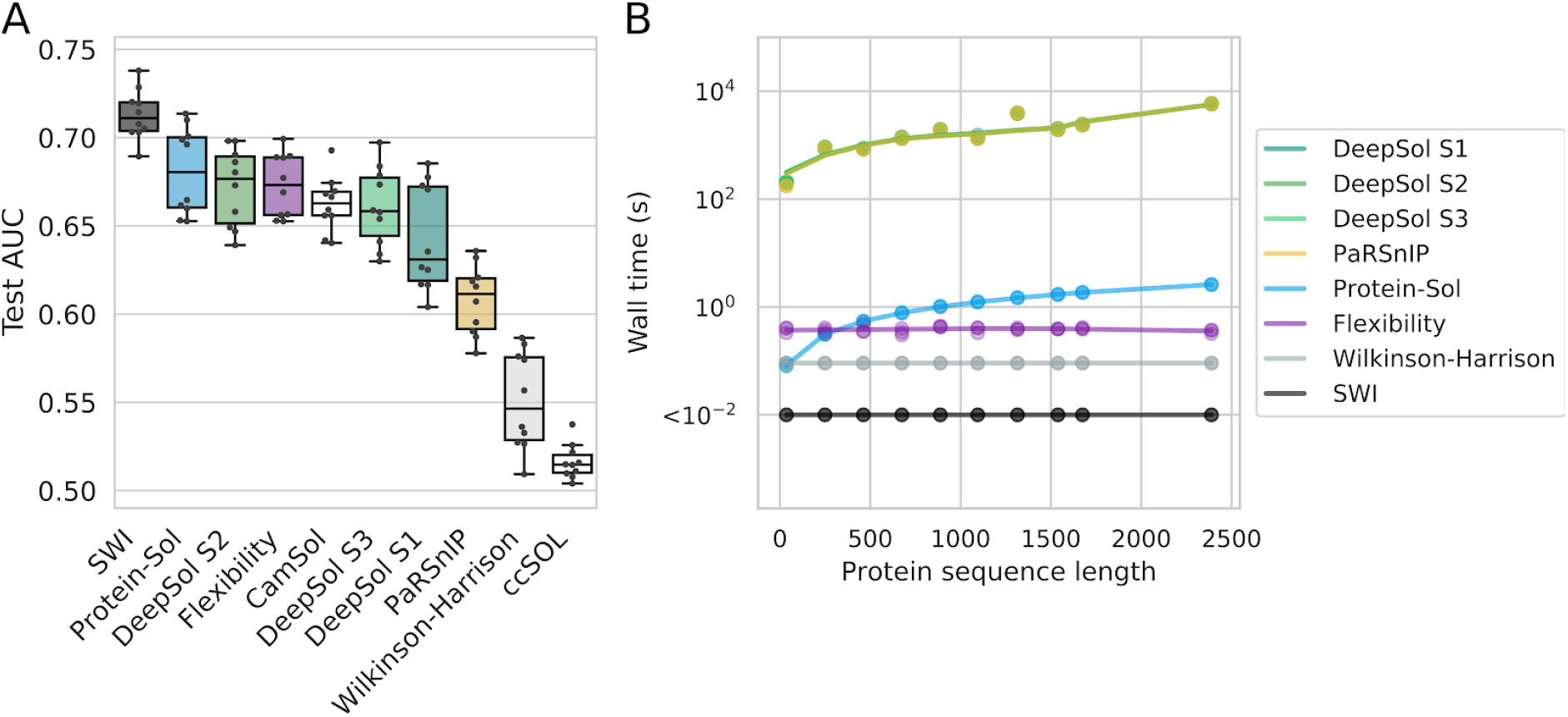
SWI outperforms existing protein solubility prediction tools. **(A)** Prediction accuracy of solubility prediction tools using the above cross-validation sets (Fig 2A). For SWI, the test AUC scores were calculated from a 10-fold cross-validation (i.e., a boxplot representation of Fig 2B). For other tools, the test AUC scores were calculated directly from the individual subsets used for cross-validation. No cross-validations were done. CamSol and ccSOL omics are only available as web servers (no fill colors). **(B)** Wall time of protein solubility prediction tools per sequence (log scale). All command line tools were run three times using 10 sequences selected from the PSI:Biology and eSOL datasets. Related data is available as Supplementary Table S7. AUC, Area Under the ROC Curve; PSI:Biology, Protein Structure Initiative:Biology; ROC, Receiver Operating Characteristic; SWI, Solubility-Weighted Index; s, seconds.

To demonstrate a use case for SWI, we developed the Soluble Domain for Protein Expression (SoDoPE) web server (see Methods and https://tisigner.com/sodope). Upon sequence submission, the SoDoPE web server enables users to navigate the protein sequence and its domains for predicting and maximising protein expression and solubility.

## DISCUSSION

Protein structural flexibility has been associated with conformal variations, functions, thermal stability, ligand binding and disordered regions (Mauno Vihinen 1987; Teague 2003; Ma 2005; Radivojac 2004; Schlessinger and Rost 2005; Yuan, Bailey, and Teasdale 2005; Yin, Li, and Li 2011). However, the use of flexibility in solubility prediction has been overlooked although their relationship has previously been noted (Tsumoto et al. 2003). In this study, we have shown that flexibility strongly correlates with solubility (Fig 3). Based on the normalised B-factors used to compute flexibility, we have derived a new position and length independent weights to score the solubility of a given protein sequence (i.e., sequence composition based score). We call this protein solubility score as SWI.

Upon further inspection, we observe some interesting properties in SWI. SWI anti-correlates with helix propensity, GRAVY, aromaticity and isoelectric point (Fig 2C and 3), suggesting that SWI incorporates the key propensities affecting solubility. Amino acid residues with a lower aromaticity or hydrophilic are known to improve protein solubility (Trevino, Martin Scholtz, and Nick Pace 2007; Niwa et al. 2009; Kramer et al. 2012; Warwicker, Charonis, and Curtis 2014; Han et al. 2019; Wilkinson and Harrison 1991). Consistent with previous studies, the charged residues aspartate (D), glutamate (E) and lysine (K) are associated with high solubility, whereas the aromatic residues phenylalanine (F), tryptophan (W) and tyrosine (Y) are associated with low solubility (Fig 2C and Supplementary Fig S7A). Cysteine residue (C) has the lowest weight probably because disulfide bonds couldn’t be properly formed in the *E. coli* expression hosts (Stewart, Aslund, and Beckwith 1998; Rosano and Ceccarelli 2014; Jia and Jeon 2016; Aslund and Beckwith 1999). The weights are likely different if the solubility analysis was done using the reductase-deficient, *E. coli* Origami host strains, or eukaryotic hosts.

Higher helix propensity has been reported to increase solubility (Idicula-Thomas and Balaji 2005; Huang et al. 2012). However, our analysis has shown that helical and turn propensities anti-correlate with solubility, whereas sheet propensity lacks correlation with solubility, suggesting that disordered regions may tend to be more soluble (Fig 3). In accordance with these, SWI has stronger negative correlations with helix and turn propensities. These findings also suggest that protein solubility can be largely explained by overall amino acid composition, not just the surface amino acid residues. This idea aligns with our understanding that protein solubility and folding are closely linked, and folding occurs cotranscriptionally, a complex process that is driven various intrinsic and extrinsic factors (Wilkinson and Harrison 1991; Chiti et al. 2003; Tartaglia et al. 2004; Diaz et al. 2010). However, it is unclear why sheet propensity has little contribution to solubility because β-sheets have been shown to link closely with protein aggregation (Idicula-Thomas and Balaji 2005).

We conclude that SWI is a well-balanced index that is derived from a simple sequence composition scoring method. To demonstrate the usefulness of SWI, we developed a web server called SoDoPE (Soluble Domain for Protein Expression; https://tisigner.com/sodope). SoDoPE calculates the probability of solubility of a user-selected region based on SWI, which can either be a full-length or a partial sequence (see Methods and Supplementary Table S8). This implementation is based on our observation that some protein domains tend to be more soluble than the others. To demonstrate this point, we have analysed three commercial monoclonal antibodies and the severe acute respiratory syndrome coronavirus proteomes (SARS-CoV and SARS-CoV-2) (Wang et al. 2009; Marra et al. 2003; F. Wu et al. 2020) (Supplementary Fig S8 and S9). These soluble domains may enhance protein solubility as a whole. SoDoPE also provides options for solubility prediction at the presence of solubility fusion tags. Similarly, solubility tags may act as soluble ‘protein domains’ that can outweigh the aggregation propensity of insoluble proteins. However, some soluble fusion proteins may become insoluble after proteolytic cleavage of solubility tags (Lebendiker and Danieli 2014). In addition, SoDoPE is integrated with TIsigner, a gene optimisation web service for protein expression. This pipeline provides a holistic approach to improve the outcome of recombinant protein expression.

## METHODS

### Protein sequence properties

The standard protein sequence properties were calculated using the Bio.SeqUtils.ProtParam module of Biopython v1.73 (Cock et al. 2009). All miscellaneous protein sequence properties were computed using the R package protr v1.6-2 (N. Xiao et al. 2015).

### Protein solubility prediction

We used the standard and miscellaneous protein sequence properties to predict the solubility of the PSI:Biology and eSOL targets (N=12,216 and 3,198, respectively) (Seiler et al. 2014; Niwa et al. 2009). For method comparison, we chose the protein solubility prediction tools that are scalable (Table 1). Default configurations were used for running the command line tools.

To benchmark the wall time of solubility prediction tools, we selected 10 sequences that span a large range of lengths from the PSI:Biology and eSOL datasets (from 36 to 2389 residues). All the tools were run and timed using a single process without using GPUs on a high performance computer [`/usr/bin/time -f ‘%E’ <command>`; CentOS Linux 7 (Core) operating system, 72 cores in 2× Broadwell nodes (E5-2695v4, 2.1 GHz, dual socket 18 cores per socket), 528 GiB memory]. Single sequence fasta files were used as input files.

### SWI

To improve protein solubility prediction, we optimised the most recently published set of normalised B-factors using the PSI:Biology dataset (Smith et al. 2003) (Fig 2). To avoid including homologous sequences in the test and training sets, we clustered the PSI:Biology targets using USEARCH v11.0.667, 32-bit (Edgar 2010). His-tag sequences were removed from all sequences before clustering to avoid false cluster inclusions. We obtained 5,050 clusters using the parameters: `-cluster_fast <input_fiie> -id 0.4 -msaout <output_fiie> -threads 4`. These clusters were divided into 10 subsets with approximately 1,200 sequences per subsets manually. The subsequent steps were done with His-tag sequences. We used Smith *et al.’* s normalised B-factors as the initial weights to maximise AUC using these 10 subsets with a 10-fold cross-validation. Since AUC is non-differentiable, we used the Nelder-Mead optimisation method (implemented in SciPy v1.2.0), which is a derivative-free, heuristic, simplex-based optimisation (Oliphant 2007; Millman and Aivazis 2011; Nelder and Mead 1965). For each step in cross-validation, we used 1,000 bootstrap resamplings containing 1,000 soluble and 1,000 insoluble proteins. Optimisation was carried out for each sample, giving 1,000 sets of weights. The arithmetic mean of these weights was used to determine the training and test AUC for the cross-validation step (Fig 2A).

### Bit score

To examine the enrichment of amino acid residues in soluble and insoluble proteins, we compute the bit scores for each amino acid residue in the PSI:Biology soluble and insoluble groups (Supplementary Fig S7A), we normalised the count of each residue (*x*) in each group by the total number of residues in that group. We used the normalised count of amino acid residues using the eSOL *E. coli* sequences as the background. The bit score of residue (*x*) for soluble or insoluble group is then given by the following equation:

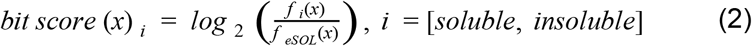

where *f_i_*(*x*) is the normalised count of residue (*x*) in the PSI:Biology soluble or insoluble group and *f_eSOL_*(*x*) is the normalised count in the eSOL sequences.

For a control, random protein sequences were generated by incrementing the length of sequence, starting from a length of 50 residues to 6,000 residues with a step size of 50 residues. A hundred random sequences were generated for each length, giving a total of 12,000 unique random sequences.

### The SoDoPE web server

To estimate the probability of solubility using SWI, we fitted the following logistic regression to the PSI:Biology dataset:

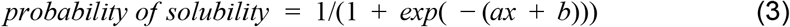

where, *x* is the SWI of a given protein sequence, *a* = 81.05812 and *b* = −62.7775. The P-value of log-likelihood ratio test was less than machine precision. Equation 3 can be used to predict the solubility of a protein sequence given that the protein is successfully expressed in *E. coli* (Supplementary Table S8).

On this basis, we developed a solubility prediction webservice called the Soluble Domain for Protein Expression (SoDoPE). Our web server accepts either a nucleotide or amino acid sequence. Upon sequence submission, a query is sent to the HMMER web server to annotate protein domains (https://www.ebi.ac.uk/Tools/hmmer/) (Potter et al. 2018). Once the protein domains are identified, users can choose a domain or any custom region (including full-length sequence) to examine the probability of solubility, flexibility and GRAVY. This functionality enables protein biochemists to plan their experiments and opt for the domains or regions with high probability of solubility. Furthermore, we implemented a simulated annealing algorithm that maximised the probability of solubility for a given region by generating a list of regions with extended boundaries. Users can also predict the improvement in solubility by selecting a commonly used solubility tag or a custom tag.

We linked SoDoPE with TIsigner, which is our existing web server for maximising the accessibility of translation initiation sites (Bhandari, Lim, and Gardner 2019). This pipeline allows users to predict and optimise both protein expression and solubility for a gene of interest. The SoDoPE web server is freely available at https://tisigner.com/sodope.

### Statistical analysis

Data analysis was done using Pandas v0.25.3 (McKinney 2010), scikit-learn v0.20.2 (Pedregosa et al. 2011), numpy v1.16.2 (van der Walt, Colbert, and Varoquaux 2011) and statsmodel v0.10.1(Seabold and Perktold 2010). Plots were generated using Matplotlib v3.0.2 (Caswell et al. 2018) and Seaborn v0.9.0 (Waskom et al. 2014).

### Code and data availability

Jupyter notebook of our analysis can be found at https://github.com/Gardner-BinfLab/SoDoPE_paper_2020. The source code for our solubility prediction server (SoDoPE) can be found at https://github.com/Gardner-BinfLab/TISIGNER-ReactJS.

## ACKNOWLEDGEMENTS

We thank New Zealand eScience Infrastructure for providing a high performance computing platform. We are grateful to Harry Biggs for proofreading our manuscript and providing feedback for the web server. This work was supported by the Ministry of Business, Innovation and Employment, New Zealand (MBIE grant: UOOX1709).

## AUTHOR CONTRIBUTIONS

C.S.L. conceived the work; B.K.B. and C.S.L. analysed the data and C.S.L. contributed flexibility analysis; B.K.B. and P.P.G formulated SWI; B.K.B. developed the SoDoPE web server; B.K.B., P.P.G. and C.S.L. wrote the manuscript.

## REFERENCES

Acton, Thomas B., Kristin C. Gunsalus, Rong Xiao, Li Chung Ma, James Aramini, Michael C. Baran, Yi-Wen Chiang, et al. 2005. “Robotic Cloning and Protein Production Platform of the Northeast Structural Genomics Consortium.” Methods in Enzymology 394: 210–43.

Agostini, Federico, Davide Cirillo, Carmen Maria Livi, Riccardo Delli Ponti, and Gian Gaetano Tartaglia. 2014. “ccSOL Omics: A Webserver for Solubility Prediction of Endogenous and Heterologous Expression in Escherichia Coli.” Bioinformatics 30 (20): 2975–77.

Aslund, F., and J. Beckwith. 1999. “The Thioredoxin Superfamily: Redundancy, Specificity, and Gray-Area Genomics.” Journal of Bacteriology 181 (5): 1375–79.

Bhandari, Bikash K., Chun Shen Lim, and Paul P. Gardner. 2019. “Highly Accessible Translation Initiation Sites Are Predictive of Successful Heterologous Protein Expression.” bioRxiv. https://doi.org/10.1101/726752.

Bhaskaran, R., and P. K. Ponnuswamy. 1988. “Positional Flexibilities of Amino Acid Residues in Globular Proteins.” International Journal of Peptide and Protein Research. https://doi.org/10.1111/j.1399-3011.1988.tb01258.x.

Caswell, Thomas A., Michael Droettboom, John Hunter, Eric Firing, Antony Lee, David Stansby, Elliott Sales de Andrade, et al. 2018. Matplotlib/matplotlib v3.0.2 (version 3.0.2). https://doi.org/10.5281/zenodo.1482099.

Chan, Wen-Ching, Po-Huang Liang, Yan-Ping Shih, Ueng-Cheng Yang, Wen-Chang Lin, and Chun-Nan Hsu. 2010. “Learning to Predict Expression Efficacy of Vectors in Recombinant Protein Production.” BMC Bioinformatics 11 Suppl 1 (January): S21.

Chen, Li, Rose Oughtred, Helen M. Berman, and John Westbrook. 2004. “TargetDB: A Target Registration Database for Structural Genomics Projects.” Bioinformatics 20 (16): 2860–62.

Chiti, Fabrizio, Massimo Stefani, Niccolò Taddei, Giampietro Ramponi, and Christopher M. Dobson. 2003. “Rationalization of the Effects of Mutations on Peptide and Protein Aggregation Rates.” Nature 424 (6950): 805–8.

Cock, Peter J. A., Tiago Antao, Jeffrey T. Chang, Brad A. Chapman, Cymon J. Cox, Andrew Dalke, Iddo Friedberg, et al. 2009. “Biopython: Freely Available Python Tools for Computational Molecular Biology and Bioinformatics.” Bioinformatics 25 (11): 1422–23.

Costa, Sofia, André Almeida, António Castro, and Lucília Domingues. 2014. “Fusion Tags for Protein Solubility, Purification and Immunogenicity in Escherichia Coli: The Novel Fh8 System.” Frontiers in Microbiology 5 (February): 63.

Craveur, Pierrick, Agnel P. Joseph, Jeremy Esque, Tarun J. Narwani, Floriane Noël, Nicolas Shinada, Matthieu Goguet, et al. 2015. “Protein Flexibility in the Light of Structural Alphabets.” Frontiers in Molecular Biosciences 2 (May): 20.

Davis, G. D., C. Elisee, D. M. Newham, and R. G. Harrison. 1999. “New Fusion Protein Systems Designed to Give Soluble Expression in Escherichia Coli.” Biotechnology and Bioengineering 65 (4): 382–88.

Diaz, Armando A., Emanuele Tomba, Reese Lennarson, Rex Richard, Miguel J. Bagajewicz, and Roger G. Harrison. 2010. “Prediction of Protein Solubility in Escherichia Coli Using Logistic Regression.” Biotechnology and Bioengineering 105 (2): 374–83.

Edgar, Robert C. 2010. “Search and Clustering Orders of Magnitude Faster than BLAST.” Bioinformatics 26 (19): 2460–61.

Esposito, Dominic, and Deb K. Chatterjee. 2006. “Enhancement of Soluble Protein Expression through the Use of Fusion Tags.” Current Opinion in Biotechnology 17 (4): 353–58.

Família, Carlos, Sarah R. Dennison, Alexandre Quintas, and David A. Phoenix. 2015. “Prediction of Peptide and Protein Propensity for Amyloid Formation.” PloS One 10 (8): e0134679.

Habibi, Narjeskhatoon, Siti Z. Mohd Hashim, Alireza Norouzi, and Mohammed Razip Samian. 2014. “A Review of Machine Learning Methods to Predict the Solubility of Overexpressed Recombinant Proteins in Escherichia Coli.” BMC Bioinformatics 15 (May): 134.

Han, Xi, Wenbo Ning, Xiaoqiang Ma, Xiaonan Wang, and Kang Zhou. 2019. “Improve Protein Solubility and Activity Based on Machine Learning Models.” bioRxiv. https://doi.org/10.1101/817890.

Harrison, R. G. 2000. “Expression of Soluble Heterologous Proteins via Fusion with NusA Protein.” Innovations 11: 4–7.

Hebditch, Max, M. Alejandro Carballo-Amador, Spyros Charonis, Robin Curtis, and Jim Warwicker. 2017. “Protein-Sol: A Web Tool for Predicting Protein Solubility from Sequence.” Bioinformatics 33 (19): 3098–3100.

Heckmann, David, Colton J. Lloyd, Nathan Mih, Yuanchi Ha, Daniel C. Zielinski, Zachary B. Haiman, Abdelmoneim Amer Desouki, Martin J. Lercher, and Bernhard O. Palsson. 2018. “Machine Learning Applied to Enzyme Turnover Numbers Reveals Protein Structural Correlates and Improves Metabolic Models.” Nature Communications 9 (1): 5252.

Hirose, Shuichi, and Tamotsu Noguchi. 2013. “ESPRESSO: A System for Estimating Protein Expression and Solubility in Protein Expression Systems.” Proteomics 13 (9): 1444–56.

Hou, Qingzhen, Raphaël Bourgeas, Fabrizio Pucci, and Marianne Rooman. 2018. “Computational Analysis of the Amino Acid Interactions That Promote or Decrease Protein Solubility.” Scientific Reports, https://doi.org/10.1038/s41598-018-32988-w.

Hou, Qingzhen, Jean-Marc Kwasigroch, Marianne Rooman, and Fabrizio Pucci. 2019. “SOLart: A Structure-Based Method to Predict Protein Solubility and Aggregation.” Bioinformatics, October. https://doi.org/10.1093/bioinformatics/btz773.

Huang, Hui-Ling, Phasit Charoenkwan, Te-Fen Kao, Hua-Chin Lee, Fang-Lin Chang, Wen-Lin Huang, Shinn-Jang Ho, Li-Sun Shu, Wen-Liang Chen, and Shinn-Ying Ho. 2012. “Prediction and Analysis of Protein Solubility Using a Novel Scoring Card Method with Dipeptide Composition.” BMC Bioinformatics 13 Suppl 17 (December): S3.

Idicula-Thomas, Susan, and Petety V. Balaji. 2005. “Understanding the Relationship between the Primary Structure of Proteins and Its Propensity to Be Soluble on Overexpression in Escherichia Coli.” Protein Science: A Publication of the Protein Society 14 (3): 582–92.

Jia, Baolei, and Che Ok Jeon. 2016. “High-Throughput Recombinant Protein Expression in Escherichia Coli: Current Status and Future Perspectives.” Open Biology 6 (8). https://doi.org/10.1098/rsob.160196.

Karplus, P. A., and G. E. Schulz. 1985. “Prediction of Chain Flexibility in Proteins.” Die Naturwissenschaften 72 (4): 212–13.

Khurana, Sameer, Reda Rawi, Khalid Kunji, Gwo-Yu Chuang, Halima Bensmail, and Raghvendra Mall. 2018. “DeepSol: A Deep Learning Framework for Sequence-Based Protein Solubility Prediction.” Bioinformatics 34 (15): 2605–13.

Kramer, Ryan M., Varad R. Shende, Nicole Motl, C. Nick Pace, and J. Martin Scholtz. 2012. “Toward a Molecular Understanding of Protein Solubility: Increased Negative Surface Charge Correlates with Increased Solubility.” Biophysical Journal. https://doi.org/10.1016/j.bpj.2012.01.060.

Kuriata, Aleksander, Valentin Iglesias, Jordi Pujols, Mateusz Kurcinski, Sebastian Kmiecik, and Salvador Ventura. 2019. “Aggrescan3D (A3D) 2.0: Prediction and Engineering of Protein Solubility.” Nucleic Acids Research 47 (W1): W300–307.

Kyte, J., and R. F. Doolittle. 1982. “A Simple Method for Displaying the Hydropathic Character of a Protein.” Journal of Molecular Biology 157 (1): 105–32.

Lebendiker, Mario, and Tsafi Danieli. 2014. “Production of Prone-to-Aggregate Proteins.” FEBS Letters 588 (2): 236–46.

Levy, E. D., S. De, and S. A. Teichmann. 2012. “Cellular Crowding Imposes Global Constraints on the Chemistry and Evolution of Proteomes.” Proceedings of the National Academy of Sciences. https://doi.org/10.1073/pnas.1209312109.

Ma, Jianpeng. 2005. “Usefulness and Limitations of Normal Mode Analysis in Modeling Dynamics of Biomolecular Complexes.” Structure 13 (3): 373–80.

Marra, Marco A., Steven J. M. Jones, Caroline R. Astell, Robert A. Holt, Angela Brooks-Wilson, Yaron S. N. Butterfield, Jaswinder Khattra, et al. 2003. “The Genome Sequence of the SARS-Associated Coronavirus.” Science 300 (5624): 1399–1404.

McKinney, Wes. 2010. “Data Structures for Statistical Computing in Python.” In Proceedings of the 9th Python in Science Conference, 51–56.

Millman, K. J., and M. Aivazis. 2011. “Python for Scientists and Engineers.” Computing in Science Engineering 13 (2): 9–12.

Natan, Eviatar, Tamaki Endoh, Liora Haim-Vilmovsky, Tilman Flock, Guilhem Chalancon, Jonathan T. S. Hopper, Bálint Kintses, et al. 2018. “Cotranslational Protein Assembly Imposes Evolutionary Constraints on Homomeric Proteins.” Nature Structural & Molecular Biology 25 (3): 279–88.

Nelder, J. A., and R. Mead. 1965. “A Simplex Method for Function Minimization.” Computer Journal 7 (4): 308–13.

Niwa, Tatsuya, Bei-Wen Ying, Katsuyo Saito, Wenzhen Jin, Shoji Takada, Takuya Ueda, and Hideki Taguchi. 2009. “Bimodal Protein Solubility Distribution Revealed by an Aggregation Analysis of the Entire Ensemble of Escherichia Coli Proteins.” Proceedings of the National Academy of Sciences of the United States of America 106 (11): 4201–6.

Oliphant, T. E. 2007. “Python for Scientific Computing.” Computing in Science Engineering 9 (3): 10–20.

Pedregosa, Fabian, Gaël Varoquaux, Alexandre Gramfort, Vincent Michel, Bertrand Thirion, Olivier Grisel, Mathieu Blondel, et al. 2011. “Scikit-Learn: Machine Learning in Python.” Journal of Machine Learning Research: JMLR 12 (Oct): 2825–30.

Potter, Simon C., Aurélien Luciani, Sean R. Eddy, Youngmi Park, Rodrigo Lopez, and Robert D. Finn. 2018. “HMMER Web Server: 2018 Update.” Nucleic Acids Research 46 (W1): W200–204.

Radivojac, P. 2004. “Protein Flexibility and Intrinsic Disorder.” Protein Science. https://doi.org/10.1110/ps.03128904.

Ragone, R., F. Facchiano, A. Facchiano, A. M. Facchiano, and G. Colonna. 1989. “Flexibility Plot of Proteins.” “Protein Engineering, Design and Selection.” https://doi.org/10.1093/protein/2.7.497.

Rawi, Reda, Raghvendra Mall, Khalid Kunji, Chen-Hsiang Shen, Peter D. Kwong, and Gwo-Yu Chuang. 2018. “PaRSnIP: Sequence-Based Protein Solubility Prediction Using Gradient Boosting Machine.” Bioinformatics. https://doi.org/10.1093/bioinformatics/btx662.

Rosano, Germán L., and Eduardo A. Ceccarelli. 2014. “Recombinant Protein Expression in Escherichia Coli: Advances and Challenges.” Frontiers in Microbiology 5 (April): 172.

Schlessinger, Avner, and Burkhard Rost. 2005. “Protein Flexibility and Rigidity Predicted from Sequence.” Proteins 61 (1): 115–26.

Seabold, Skipper, and Josef Perktold. 2010. “Statsmodels: Econometric and Statistical Modeling with Python.” In Proceedings of the 9th Python in Science Conference. http://conference.scipy.org/proceedings/scipy2010/pdfs/seabold.pdf.

Seiler, Catherine Y., Jin G. Park, Amit Sharma, Preston Hunter, Padmini Surapaneni, Casey Sedillo, James Field, et al. 2014. “DNASU Plasmid and PSI:Biology-Materials Repositories: Resources to Accelerate Biological Research.” Nucleic Acids Research 42 (Database issue): D1253–60.

Smith, David K., Predrag Radivojac, Zoran Obradovic, A. Keith Dunker, and Guang Zhu. 2003. “Improved Amino Acid Flexibility Parameters.” Protein Science: A Publication of the Protein Society 12 (5): 1060–72.

Sormanni, Pietro, Leanne Amery, Sofia Ekizoglou, Michele Vendruscolo, and Bojana Popovic. 2017. “Rapid and Accurate in Silico Solubility Screening of a Monoclonal Antibody Library.” Scientific Reports 7 (1): 8200.

Sormanni, Pietro, Francesco A. Aprile, and Michele Vendruscolo. 2015. “The CamSol Method of Rational Design of Protein Mutants with Enhanced Solubility.” Journal of Molecular Biology, https://doi.org/10.1016/j.jmb.2014.09.026.

Stewart, E. J., F. Aslund, and J. Beckwith. 1998. “Disulfide Bond Formation in the Escherichia Coli Cytoplasm: An in Vivo Role Reversal for the Thioredoxins.” The EMBO Journal 17 (19): 5543–50.

Tartaglia, Gian Gaetano, Andrea Cavalli, Riccardo Pellarin, and Amedeo Caflisch. 2004. “The Role of Aromaticity, Exposed Surface, and Dipole Moment in Determining Protein Aggregation Rates.” Protein Science: A Publication of the Protein Society 13 (7): 1939.

Teague, Simon J. 2003. “Implications of Protein Flexibility for Drug Discovery.” Nature Reviews. Drug Discovery 2 (7): 527–41.

Trevino, Saul R., J. Martin Scholtz, and C. Nick Pace. 2007. “Amino Acid Contribution to Protein Solubility: Asp, Glu, and Ser Contribute More Favorably than the Other Hydrophilic Amino Acids in RNase Sa.” Journal of Molecular Biology, https://doi.org/10.1016/j.jmb.2006.10.026.

Tsumoto, Kouhei, Daisuke Ejima, Izumi Kumagai, and Tsutomu Arakawa. 2003. “Practical Considerations in Refolding Proteins from Inclusion Bodies.” Protein Expression and Purification 28 (1): 1–8.

Vihinen, Mauno. 1987. “Relationship of Protein Flexibility to Thermostability.” “Protein Engineering, Design and Selection.” https://doi.org/10.1093/protein/1.6.477.

Vihinen, M., E. Torkkila, and P. Riikonen. 1994. “Accuracy of Protein Flexibility Predictions.” Proteins 19 (2): 141–49.

Waldo, Geoffrey S. 2003. “Genetic Screens and Directed Evolution for Protein Solubility.” Current Opinion in Chemical Biology 7 (1): 33–38.

Walt, Stéfan van der, S. Chris Colbert, and Gaël Varoquaux. 2011. “The NumPy Array: A Structure for Efficient Numerical Computation.” Computing in Science & Engineering 13 (2): 22–30.

Wang, Xiaoling, Tapan K. Das, Satish K. Singh, and Sandeep Kumar. 2009. “Potential Aggregation Prone Regions in Biotherapeutics: A Survey of Commercial Monoclonal Antibodies.” mAbs 1 (3): 254–67.

Warwicker, Jim, Spyros Charonis, and Robin A. Curtis. 2014. “Lysine and Arginine Content of Proteins: Computational Analysis Suggests a New Tool for Solubility Design.” Molecular Pharmaceutics 11 (1): 294–303.

Waskom, Michael, Olga Botvinnik, Paul Hobson, John B. Cole, Yaroslav Halchenko, Stephan Hoyer, Alistair Miles, et al. 2014. “Seaborn: v0.5.0 (November 2014),” November. https://doi.org/10.5281/zenodo.12710.

Wilkinson, D. L., and R. G. Harrison. 1991. “Predicting the Solubility of Recombinant Proteins in Escherichia Coli.” Bio/technology 9 (5): 443–48.

Wu, Fan, Su Zhao, Bin Yu, Yan-Mei Chen, Wen Wang, Yi Hu, Zhi-Gang Song, et al. 2020. “Complete Genome Characterisation of a Novel Coronavirus Associated with Severe Human Respiratory Disease in Wuhan, China.” bioRxiv, https://doi.org/10.1101/2020.01.24.919183.

Wu, Zachary, S. B. Jennifer Kan, Russell D. Lewis, Bruce J. Wittmann, and Frances H. Arnold. 2019. “Machine Learning-Assisted Directed Protein Evolution with Combinatorial Libraries.” Proceedings of the National Academy of Sciences of the United States of America 116 (18): 8852–58.

Xiao, Nan, Dong-Sheng Cao, Min-Feng Zhu, and Qing-Song Xu. 2015. “protr/ProtrWeb: R Package and Web Server for Generating Various Numerical Representation Schemes of Protein Sequences.” Bioinformatics 31 (11): 1857–59.

Xiao, Rong, Stephen Anderson, James Aramini, Rachel Belote, William A. Buchwald, Colleen Ciccosanti, Ken Conover, et al. 2010. “The High-Throughput Protein Sample Production Platform of the Northeast Structural Genomics Consortium.” Journal of Structural Biology 172 (1): 21–33.

Yang, Kevin K., Zachary Wu, and Frances H. Arnold. 2019. “Machine-Learning-Guided Directed Evolution for Protein Engineering.” Nature Methods 16 (8): 687–94.

Yin, Hui, Yi-Zhou Li, and Meng-Long Li. 2011. “On the Relation between Residue Flexibility and Residue Interactions in Proteins.” Protein and Peptide Letters 18 (5): 450–56.

Yuan, Zheng, Timothy L. Bailey, and Rohan D. Teasdale. 2005. “Prediction of Protein B-Factor Profiles.” Proteins: Structure, Function, and Bioinformatics. https://doi.org/10.1002/prot.20375.

