## Supplementary Fig for "Solubility-Weighted Index: fast and accurate prediction of protein solubility"

### SUPPLEMENTARY NOTES

The B-factor or temperature factor of the atom in a crystalline structure is the measure of mean squared displacement $(u = \langle (x - x_0)^2 \rangle)$, where $x$ is the displacement of the atom from its mean position $x_0$. B-factor thus reflects the orderedness of the crystal lattice and subsequent uncertainty in X-ray scattering structure determination (Schlessinger and Rost, 2005; Carugo, 2018; Bramer and Wei, 2018).

$$B = 8\pi^2 u \qquad \text{(i)}$$

Since the distribution of B-factors varies with protein crystal structures, experimentally determined B-factors (for example from the Protein Data Bank) are not generalisable without appropriate normalisation. To address this issue, the B-factors of $C_\alpha$ atoms are collected from a number of high-resolution protein structures and normalised. The normalisation is often done by Z scoring, for example, for a residue $i$, $B^i_{norm} = (B^i - \langle B \rangle)/\sigma$, where $\sigma$ is the standard deviation and $\langle B \rangle$ is the mean of B-factors within the polypeptide chain (Schlessinger and Rost, 2005; Smith *et al.*, 2003; Karplus and Schulz, 1985; Vihinen *et al.*, 1994).

The profile of normalised B-factors along a protein sequence can be calculated using a sliding window approach [e.g., 9 amino acid residues as implemented in Biopython (Vihinen *et al.*, 1994; Cock *et al.*, 2009)]. The profile plot can be used to visualise and infer the local flexibility and dynamics of the protein structure (Karplus and Schulz, 1985; Vihinen *et al.*, 1994). Previous studies that formulated flexibility also compared their computed values with the B-factors of previously solved protein structures using correlation tests (Vihinen, 1987; Vihinen *et al.*, 1994).

To calculate global structural flexibility, we reasoned that Vihinen *et al.*'s sliding window method can be approximated by a more straightforward arithmetic mean. This sliding window method computes the local flexibility $f_i$ of a given amino acid residue $i$ as:

$$\begin{aligned} f_i = \tfrac{1}{5.25}[B_i &+ 0.8125(B_{i-1} + B_{i+1}) + 0.625(B_{i-2} + B_{i+2}) \\ &+ 0.4375(B_{i-3} + B_{i+3}) + 0.25(B_{i-4} + B_{i+4}) \end{aligned} \qquad \text{(ii)}$$

where $B_i$ is the normalised B-factor of the $i^{th}$ $C_\alpha$ atom and so on. The arithmetic mean of these $f_i$ can be approximately written as:

$$\begin{aligned} \langle f_i \rangle &\approx \tfrac{1}{5.25(n-9)}(1 + 2 \times (0.8125 + 0.625 + 0.4375 + 0.25)) \sum_{i=5}^{n-4} B_i \\ &= \tfrac{1}{(n-9)} \sum_{i=5}^{n-4} B_i \end{aligned} \qquad \text{(iii)}$$

where $n$ is the number of residues in the protein. For sequence composition scoring, the arithmetic mean of $B_i$ of a given full-length sequence is written as:

$$< B > \; = \; \frac{1}{n}\left(\sum_{i=1}^{n} B_i\right) \qquad \text{(iv)}$$

Dividing (iii) by (iv), and approximating that the sums run at equal intervals, we can write:

$$\frac{<f_i>}{<B>} \approx \frac{n}{(n-9)} \qquad \text{(v)}$$

$\frac{n}{(n-9)}$ is monotonically decreasing for $n \geq 10$ and quickly approaches 1 with an increasing $n$. Thus, $< f_i >$ is nearly equal to $< B >$ and they are strongly correlated.

### SUPPLEMENTARY FIGURES

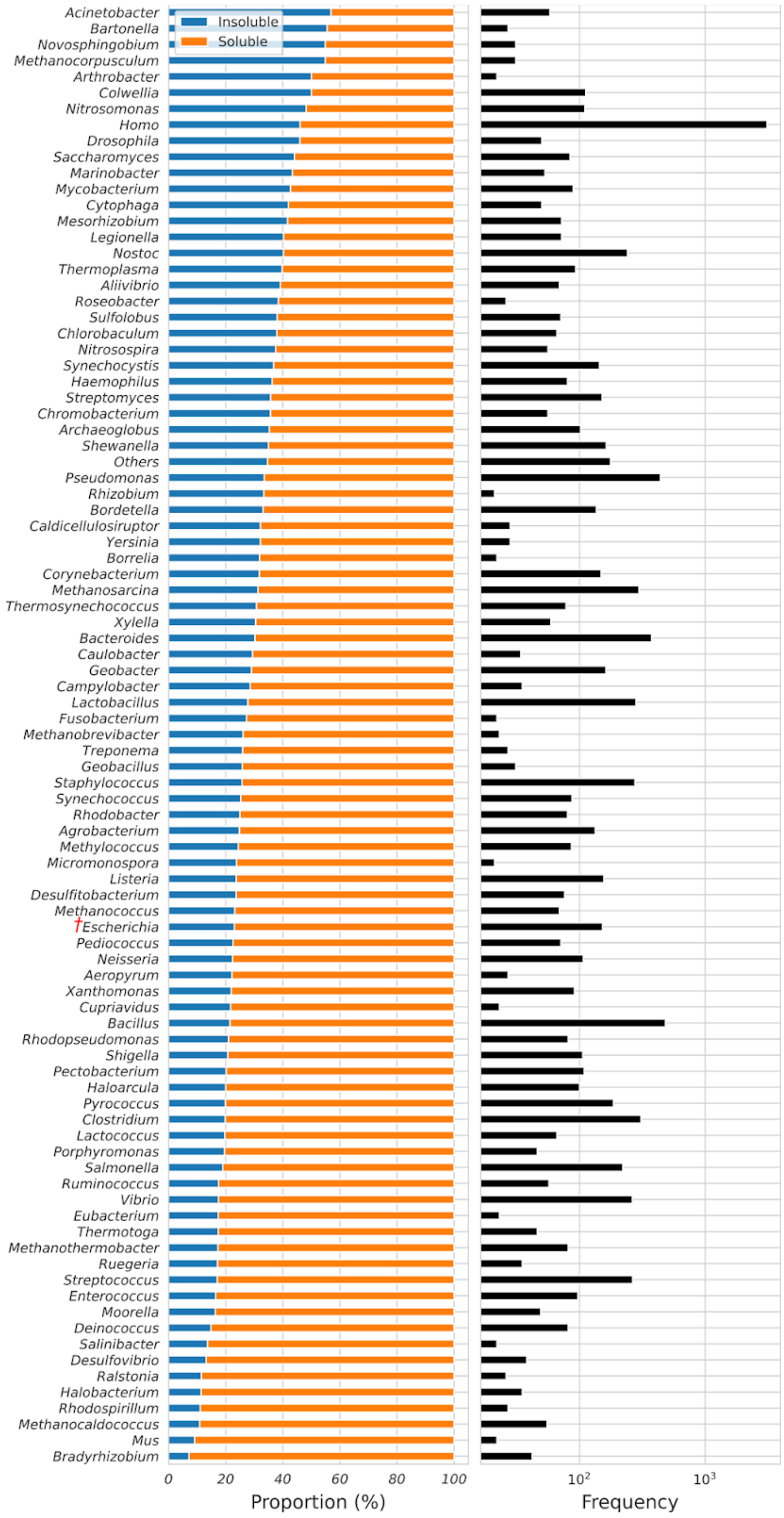


**Fig S1.** Solubility of the PSI:Biology targets grouped by source. A total of 12,216 PSI:Biology targets from over 196 species were analysed in this study (8,238 soluble and 3,978 insoluble proteins). Genera with at least 20 target genes are shown and the remaining as 'Others'. Red obelisk indicates *E. coli*.

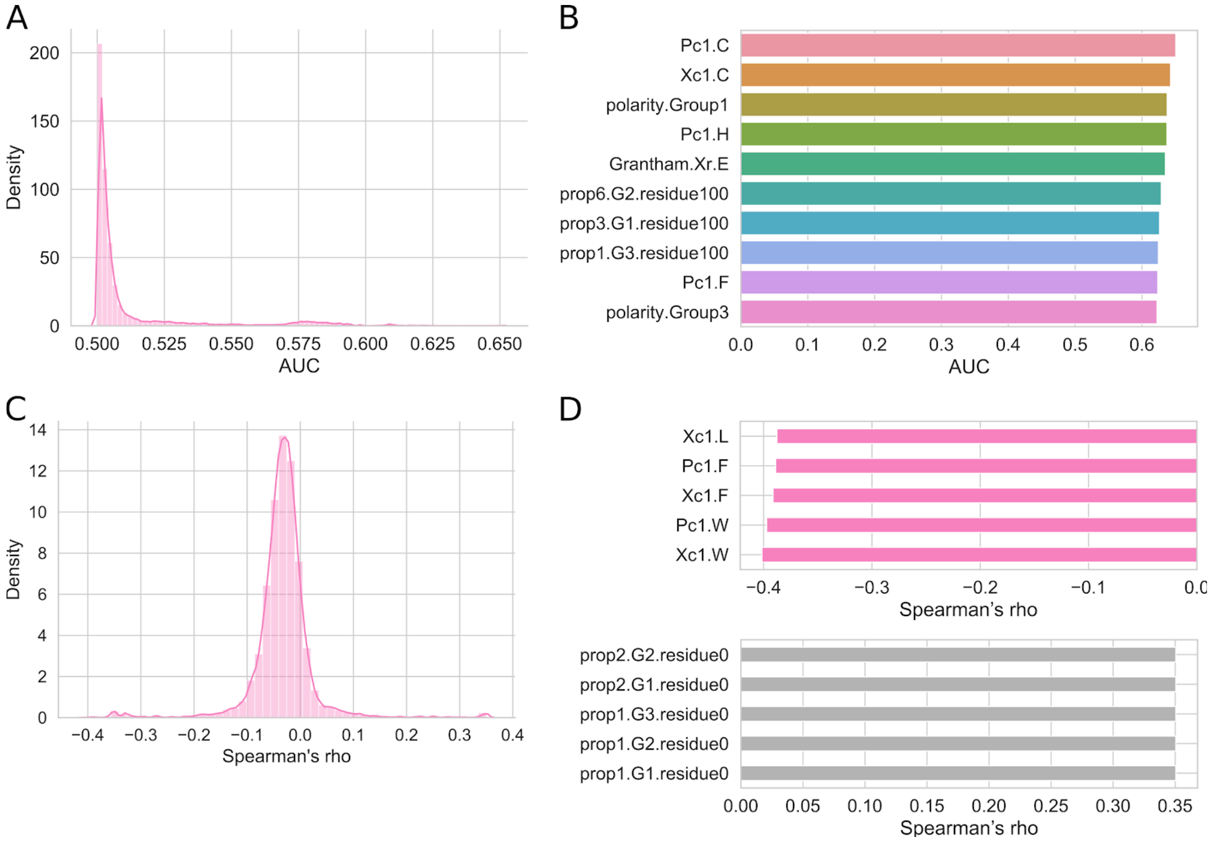


**Fig S2. Prediction accuracy of 9,920 miscellaneous protein sequence properties. (A)** Density distribution of AUC scores shows that relatively few features have high prediction accuracy (PSI:Biology dataset, N = 12,216). **(B)** Top-ranked features by AUC scores, which include the (amphiphilic) pseudo-amino acid compositions for cysteine residues (Pc1.C and Xc.1.C). **(C)** Density distribution of Spearman's rho shows that relatively few features have strong correlation coefficients with *E. coli* protein solubility (eSOL dataset, N = 3,198). **(D)** Top-ranked features by Spearman's correlation coefficients, which include the (amphiphilic) pseudo-amino acid compositions for aromatic amino acid residues (Xc1.W, Pc1.W, Xc1.F, and Pc1.F). The complete list of AUC scores and Spearman's correlation coefficients are available in Supplementary Table S2. AUC, Area Under the ROC Curve; Pc1, amphiphilic pseudo-amino acid composition; polarity.Group1, one of the three groups of amino acid residues based on polarity (L, I, F, W, C, M, V, Y); polarity.Group3, one of the three groups of amino acid residues based on polarity (H, Q, R, K, N, E, D); prop{1-7}.G{1,2,3}.residue{0,25,50,100%}, position percent for one of the three groups of amino acid residues by one of the seven properties listed in Table 1 of the protr vignettes, https://cran.r-project.org/web/packages/protr/vignettes/protr.html; PSI:Biology, Protein Structure Initiative:Biology; ROC, Grantham.Xr, Quasi-sequence-order based on Grantham's chemical distance matrix; Receiver Operating Characteristic; Xc1, pseudo-amino acid composition.

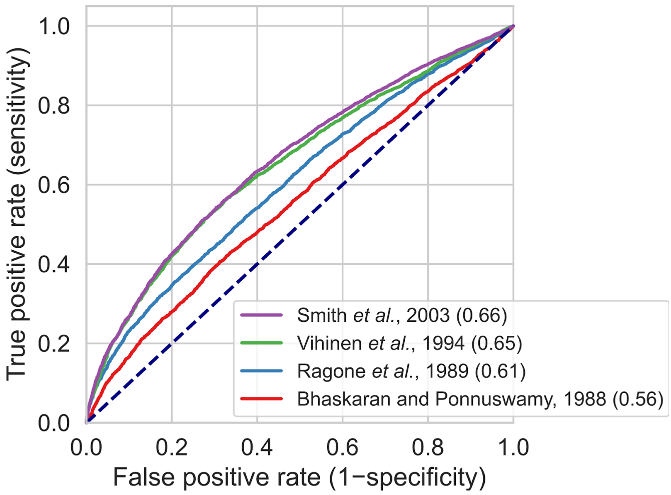


**Fig S3. ROC analysis of sequence composition scores for solubility using previously published sets of normalised B-factors.** The PSI:Biology dataset (N = 12,216) was used for solubility prediction. AUC scores (perfect = 1.00, random = 0.50) are shown in parentheses. Dashed lines denote the performance of random classifiers. PSI:Biology, Protein Structure Initiative:Biology; ROC, Receiver Operating Characteristic.

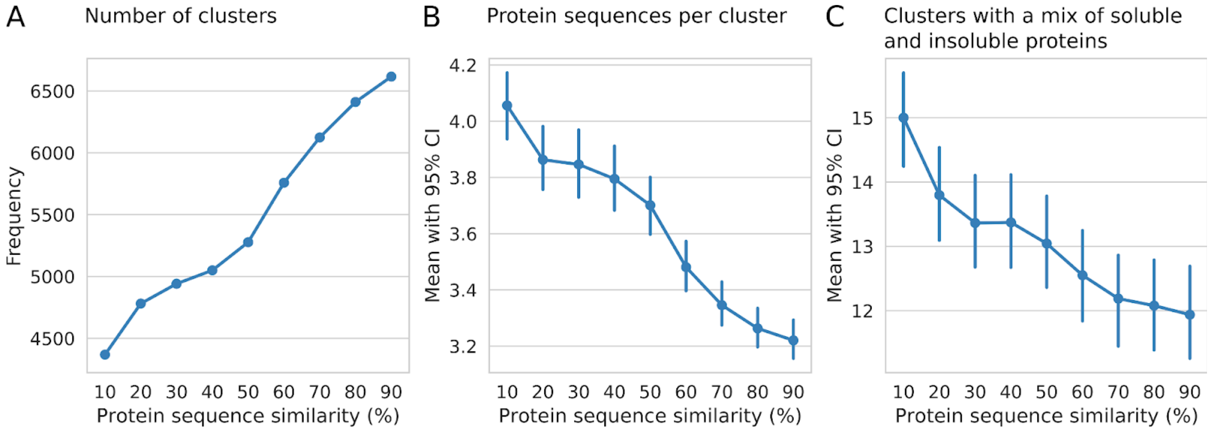


**Fig S4. Relationship between protein solubility and sequence similarity, related to Fig 2.** USEARCH was used to cluster the PSI:Biology targets (N = 12,216) at different percent similarity cutoffs (using the parameter `-id 0.1` to `0.9`; see https://drive5.com/usearch/manual/uclust_algo.html). **(A)** High numbers of clusters across different similarity cutoffs and **(B)** low numbers of sequences per cluster indicate that the PSI:Biology targets are highly diverse (Supplementary Fig S1). **(C)** Over about 12% of clusters contain a mix of soluble and insoluble proteins across different similarity cutoffs. CI, Confidence Intervals.

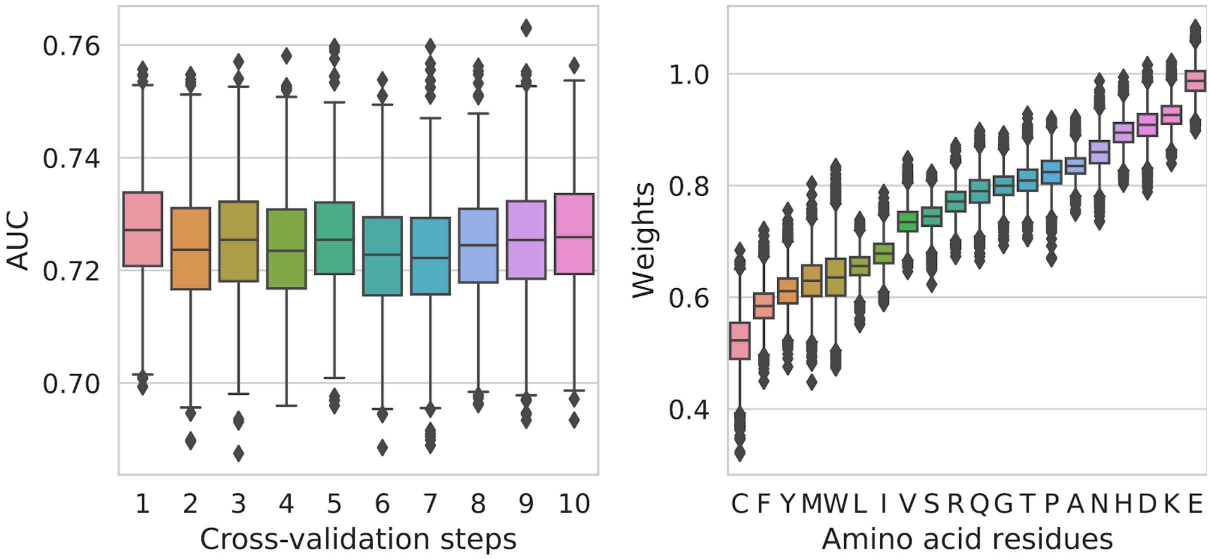


**Fig S5. AUC scores and weights of amino acid residues obtained from individual bootstrap samples, related to Fig 2.** For each cross-validation step, 1,000 soluble and 1,000 insoluble proteins were resampled 1,000 times. For each bootstrap resampling, the weights of amino acid residues were optimised by maximising AUC using the Nelder-Mead algorithm. The optimised weights, i.e., the arithmetic means of the weights of individual amino acid residues in each cross-validation step, were used for sequence composition scoring. The training and test AUC scores were subsequently calculated (Fig 2B, 4A and Supplementary Table S3). AUC, Area Under the ROC Curve; ROC, Receiver Operating Characteristic.

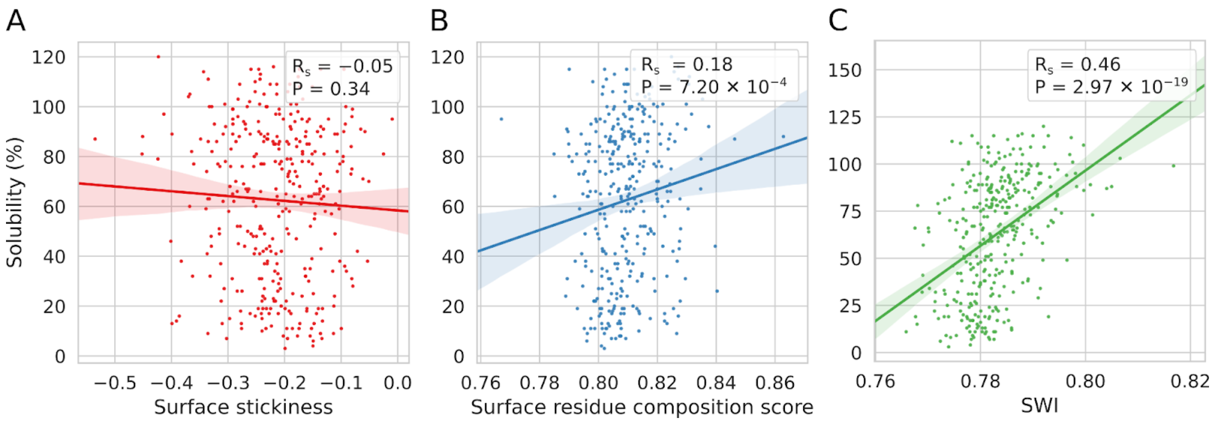


**Fig S6. Relationship between protein solubility and surface amino acid residues.** The analyses were done using eSOL and the surface 'stickiness' of *E. coli* proteins (N = 348). **(A)** Protein solubility has a low correlation with surface 'stickiness'. **(B)** A low correlation was obtained after maximising the correlation between solubility and the surface residue composition scores using the Nelder-Mead algorithm. Smith *et al.*'s normalised B-factors were used as initial weights. **(C)** In contrast, protein solubility has a stronger correlation with SWI. $R_s$, Spearman's rho; SWI, Solubility-Weighted Index.

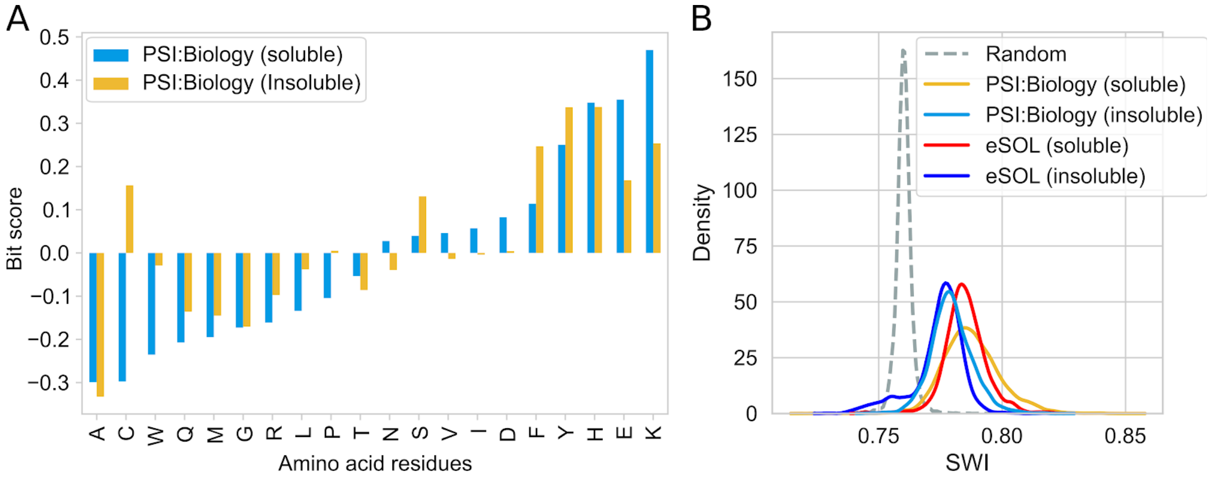


**Fig S7. Properties of soluble and insoluble proteins. (A)** Enrichment of amino acid residues in the PSI:Biology targets relative to the eSOL sequences (N = 12,216 and 3,198, respectively). **(B)** Distribution of the SWI for soluble and insoluble proteins, and random sequences. The eSOL sequences were grouped into soluble and insoluble proteins, i.e, <30% and >70% solubility cutoffs, respectively (Supplementary Table S1B). Random sequences were generated from a length of 50 to 6,000 amino acid residues, with an increment of 50 residues. A total of 12,000 random sequences were generated, 100 sequences for each length. PSI:Biology, Protein Structure Initiative:Biology; SWI, Solubility-Weighted Index.

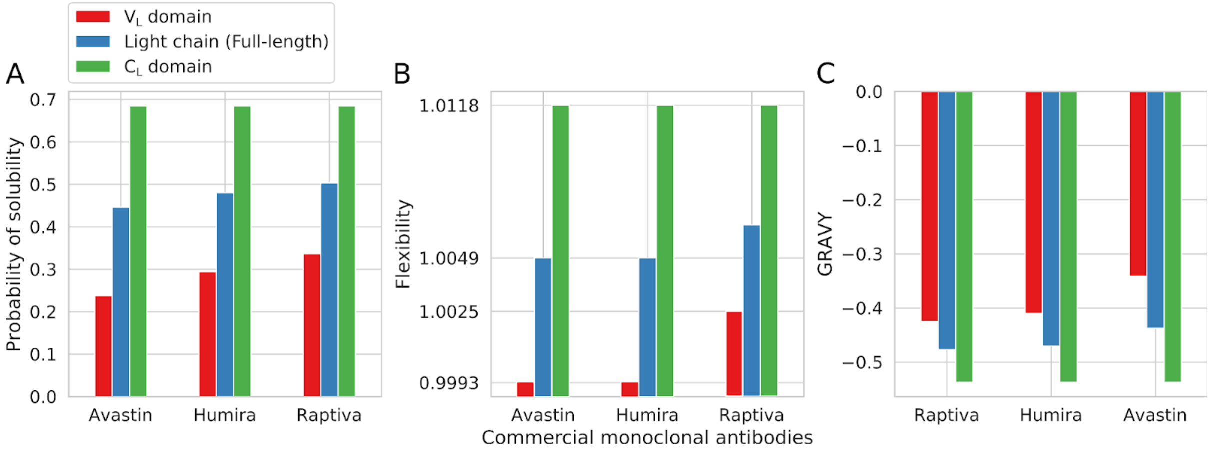


**Fig S8. Solubility analysis of three commercial monoclonal antibodies.** The variable domains of immunoglobulin light chains ($V_L$) have **(A)** lower probabilities of solubility, **(B)** lower structural flexibilities (log scale), and **(C)** higher GRAVY than the constant domains ($C_L$). The sequences of Avastin (216974-75-3), Humira (331731-18-1), and Raptiva (214745-43-4) were retrieved from the Common Chemistry database. CAS registry numbers are shown in parentheses. GRAVY, Grand Average of Hydropathy.

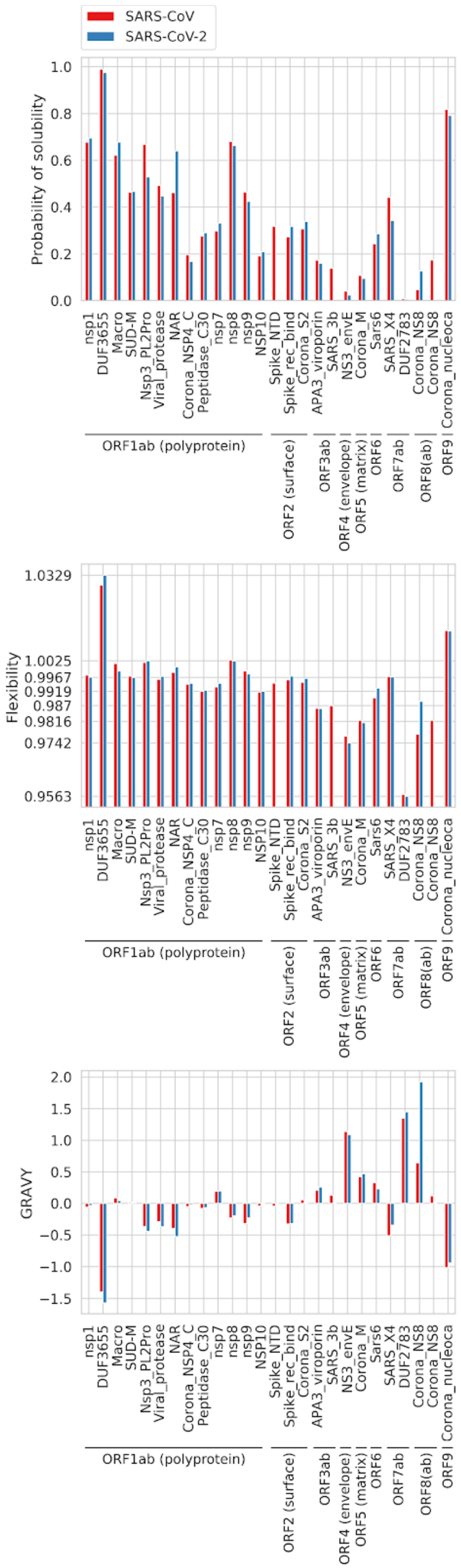


**Fig S9. Solubility analysis of the SARS-CoV and SARS-CoV-2 proteomes**. The viral proteomes were retrieved from NCBI RefSeq on 23 March 2020 (NC_004718.3 and NC_045512.2). The polypeptides/domains were annotated by the HMMER web server using the Pfam database. No domains were annotated for ORF10. The ORF2, 4, 5, and 8b proteins/domains have low probabilities of solubility, whereas the ORF9 protein have a high probability of solubility, which are consistent with previous protein expression studies (Wu *et al.*, 2004; Kam *et al.*, 2007; Neuman *et al.*, 2011; Shi *et al.*, 2019). The flexibility plot is shown in log scale. GRAVY, Grand Average of Hydropathy; SARS-CoV, severe acute respiratory syndrome coronavirus; SARS-CoV-2, severe acute respiratory syndrome coronavirus 2.

### REFERENCES


Bramer,D. and Wei,G.-W. (2018) Blind prediction of protein B-factor and flexibility. *J. Chem. Phys.*, **149**, 134107.

Carugo,O. (2018) How large B-factors can be in protein crystal structures. *BMC Bioinformatics*, **19**, 61.

Cock,P.J.A. *et al.* (2009) Biopython: freely available Python tools for computational molecular biology and bioinformatics. *Bioinformatics*, **25**, 1422–1423.

Kam,Y.W. *et al.* (2007) Antibodies against trimeric S glycoprotein protect hamsters against SARS-CoV challenge despite their capacity to mediate FcgammaRII-dependent entry into B cells in vitro. *Vaccine*, **25**, 729–740.

Karplus,P.A. and Schulz,G.E. (1985) Prediction of chain flexibility in proteins. *Naturwissenschaften*, **72**, 212–213.

Neuman,B.W. *et al.* (2011) A structural analysis of M protein in coronavirus assembly and morphology. *J. Struct. Biol.*, **174**, 11–22.

Schlessinger,A. and Rost,B. (2005) Protein flexibility and rigidity predicted from sequence. *Proteins*, **61**, 115–126.

Shi,C.-S. *et al.* (2019) SARS-Coronavirus Open Reading Frame-8b triggers intracellular stress pathways and activates NLRP3 inflammasomes. *Cell Death Discovery*, **5**, 1–12.

Smith,D.K. *et al.* (2003) Improved amino acid flexibility parameters. *Protein Sci.*, **12**, 1060–1072.

Vihinen,M. *et al.* (1994) Accuracy of protein flexibility predictions. *Proteins*, **19**, 141–149.

Vihinen,M. (1987) Relationship of protein flexibility to thermostability. *'Protein Engineering, Design and Selection'*, **1**, 477–480.

Wu,X.D. *et al.* (2004) The spike protein of severe acute respiratory syndrome (SARS) is cleaved in virus infected Vero-E6 cells. *Cell Res.*, **14**, 400–406.
